# Multivariate GWAS elucidates the genetic architecture of alcohol consumption and misuse, corrects biases, and reveals novel associations with disease

**DOI:** 10.1101/2020.09.21.304196

**Authors:** Travis T Mallard, Jeanne E Savage, Emma C Johnson, Yuye Huang, Alexis C Edwards, Jouke J Hottenga, Andrew D Grotzinger, Daniel E Gustavson, Mariela V Jennings, Andrey Anokhin, Danielle M Dick, Howard J Edenberg, John R Kramer, Dongbing Lai, Jacquelyn L Meyers, Ashwini K Pandey, Kathryn Paige Harden, Michel G Nivard, Eco JC de Geus, Dorret I Boomsma, Arpana Agrawal, Lea K Davis, Toni-Kim Clarke, Abraham A Palmer, Sandra Sanchez-Roige

**Author notes:** Corresponding author: Sandra Sanchez-Roige.

## Abstract

Genome-wide association studies (GWASs) of the Alcohol Use Disorder Identification Test (AUDIT), a ten-item screener for alcohol use disorder (AUD), have elucidated novel loci for alcohol consumption and misuse. However, these studies also revealed that GWASs can be influenced by numerous biases (e.g., measurement error, selection bias), which have led to inconsistent genetic correlations between alcohol involvement and AUD, as well as paradoxically negative genetic correlations between alcohol involvement and psychiatric disorders/medical conditions. To explore these unexpected differences in genetic correlations, we conducted the first item-level and largest GWAS of AUDIT items (N=160,824), and applied a multivariate framework to mitigate previous biases. In doing so, we identified novel patterns of similarity (and dissimilarity) among the AUDIT items, and found evidence of a correlated two-factor structure at the genetic level (Consumption and Problems, rg=.80). Moreover, by applying empirically-derived weights to each of the AUDIT items, we constructed an aggregate measure of alcohol consumption that is strongly associated with alcohol dependence (rg=.67) and several other psychiatric disorders, and no longer positively associated with health and positive socioeconomic outcomes. Lastly, by performing polygenic analyses in three independent cohorts that differed in their ascertainment and prevalence of AUD, we identified novel genetic associations between alcohol consumption, alcohol misuse, and human health. Our work further emphasizes the value of AUDIT for both clinical and genetic studies of AUD, and the importance of using multivariate methods to study genetic associations that are more closely related to AUD.

## INTRODUCTION

Over the past decade, genome-wide association studies (**GWASs**) have advanced our understanding of alcohol use disorders (**AUDs**)(1). Many of these studies have relied on a categorical classification, comparing clinically-ascertained AUD cases and controls (e.g., 2), whereas others have employed a complementary approach using dimensional measures of alcohol consumption and screener-based AUD symptoms in population-based cohorts (e.g., 3–6). Often, these dimensional measures can more easily be administered at scale via self-report questionnaires than can clinical diagnostic measures, thereby accelerating genetic discovery through drastic increases in sample size. The Alcohol Use Disorders Identification Test (**AUDIT**)(7), a ten-item questionnaire that screens for drinking habits and problems by measuring aspects of alcohol use and misuse in the past year, is one such measure. The AUDIT differentiates between two related but distinct facets of AUD: alcohol consumption (sum of items 1-3, “**AUDIT-C**”), which is necessary but not sufficient for a diagnosis of AUD, and problematic consequences of alcohol consumption (sum of items 4-10, “**AUDIT-P**”), which more closely resemble the diagnostic criteria of AUD. A recent meta-analysis of AUD and AUDIT-P GWASs identified 29 novel loci (5), representing one of the biggest advances of AUD genetics to date (2–4, 6).

Several studies using self-report instruments have now revealed that not all aspects of alcohol involvement are interchangeable. While AUDIT can be used as a unidimensional screener (i.e., AUDIT-Total, summing both AUDIT-C and AUDIT-P items), previous research has shown that AUDIT can differentiate between two related but distinct facets of AUD. For example, we previously showed that AUDIT-C and AUDIT-P have distinct genetic relationships with clinically-defined AUD (6), as well as other forms of psychopathology. Surprisingly, AUDIT-C was *only moderately* positively genetically associated with alcohol dependence, *positively* associated with socioeconomic variables, and *negatively* associated with some forms of psychopathology, whereas AUDIT-P showed strong positive associations with alcohol dependence and numerous other psychiatric disorders.

Although this divergence might reflect true differences in the biology underlying alcohol consumption versus problems, it may be confounded by other factors, such as sources of selection bias, genetic heterogeneity among the individual items, and measurement error (1, 8). Furthermore, AUDIT-C and AUDIT-P are constructed using a composite score approach that assumes (i) the scale is unidimensional, and (ii) each item is equally informative of the construct being measured. This approach is not based on any empirical property of the data – it is a holdover from its original intent as a screener for primary health care settings – and the resulting weights potentially introduce error in downstream analyses. While these issues have been thoroughly studied at the phenotypic level via factor analysis (**Table S1**), they have not yet been investigated at the genetic level. Using methods that can account for or mitigate such measurement problems will allow researchers to capitalize on the potential of dimensional measures like AUDIT for genetic discovery.

In the present study, we sought to elucidate the genetics of alcohol consumption and problematic consequences of alcohol use measured via AUDIT using a novel multivariate framework. We performed the first item-level and largest to-date GWAS meta-analyses of AUDIT (N=160,824), using data from three population-based cohorts. Next, we used Genomic Structural Equation Modelling (**Genomic SEM**)(9) to investigate the latent genetic structure of the ten AUDIT items based on prior knowledge (**Table S1**), and perform multivariate GWASs of alcohol consumption (items 1-3) and problematic use (items 4-10). Crucially, this approach allowed us to apply more nuanced, empirically-derived weights to each of the AUDIT items when constructing our aggregate measures (as opposed to giving each item equivalent weight), which is a novel approach for GWASs of AUD phenotypes. Finally, to characterize the biology and liability associated with each latent genetic factor, we used a variety of *in-silico* tools and polygenic analyses spanning three independent cohorts that varied in their method of ascertainment and prevalence of AUD.

We hypothesized that a higher resolution of each of the alcohol phenotypes measured in AUDIT would further our understanding of the differences among indices of alcohol consumption [*e*.*g*., frequency (item 1), quantity (item 2), binge drinking (item 3)] and problematic alcohol use, and how they relate to human health. We anticipated that the genetic contributions to alcohol consumption and problematic use would not be completely overlapping, and that genomic modeling using item-level data would ameliorate the confounding associations between alcohol consumption, AUD, and indices of health that complicated previous GWAS efforts.

## MATERIALS AND METHODS

### Discovery samples and phenotype construction

We collected AUDIT (7) and genotype data from three population-based cohorts: UK Biobank (**UKB**, *n*_max_=147,267), the Netherlands Twin Register (**NTR**, *n*_max_=9,975), and the Avon Longitudinal Study of Parents and Children (**ALSPAC**, *n*_max_=3,582). We used the same phenotyping strategies across the three cohorts, which are described in the **Supplementary Material 2**. AUDIT scores and demographics for each cohort are reported in **Table S2**. Genotyping, imputation and quality control (**QC**) procedures have been extensively described in previous publications (10–12). Because AUDIT was administered with skip logic in UKB, we used multiple imputation by chained equations to minimize the impact of missing data on our item-level GWAS (see **Supplementary Material 2**.**1** for details).

### Univariate genome-wide association and meta-analyses

In UKB, we used BOLT-LMM (13) v2.3.2 to perform GWASs for each of the ten AUDIT items, with the first 40 ancestry principal components (**PCs**), sex, age, sex-by-age interactions, and batch as covariates. In NTR, we used the fastgwa function of GCTA (14) to perform GWASs using the first 5 ancestry PCs, sex, birth year, and genotyping platform as covariates. In ALSPAC, we used PLINK v2.0 (15) to conduct GWAS after excluding related individuals, with the first 10 ancestry PCs, sex, and age as covariates. Note that both BOLT-LMM and fastgwa are capable of analyzing related individuals. Further details are included in the **Supplementary Material 3** and prior work (16). We then used METAL (17) to perform sample-size weighted meta-analyses of the cohort-level GWAS summary statistics for each AUDIT item following QC procedures (see **Supplementary Material 4**). A total of 8,596,116 SNPs were included in the meta-analyses.

### Phenotypic and genetic correlations

We used the lavaan (18) v0.6.5 package in R to estimate polychoric phenotypic correlations (*r*_p_) among AUDIT items. We used the Genomic SEM v0.0.2 package in R, which is based on LD score regression (**LDSC**)(19), to estimate the heritability of each of the ten AUDIT items, and the genetic correlations between them. We applied standard QC procedures (e.g., used precomputed LD scores, excluded the major histocompatibility region, SNPs restricted to HapMap 3, applied MAF ≥1% and information score >.90 filters). Lastly, we used Genomic SEM (9) to estimate genetic correlations between latent genetic factors and complex traits and disorders broadly related to human health. We applied a standard Benjamini–Hochberg false discovery rate (**FDR** 5%) correction to account for multiple testing.

### Phenotypic and genetic factor analysis

We used the lavaan (18) v0.6.5 and Genomic SEM (9) v0.0.2 packages in R to conduct phenotypic and genetic confirmatory factor analyses, respectively, using weighted least squares estimation. We tested three models: (i) a parallel factor model (i.e., a sum score model), (ii) a common factor model, and (iii) a correlated factors model. The common and correlated-factors models were selected based on prior research (**Table S1**) while the parallel factor model served to test the restrictive assumptions of sum score approaches. We assessed model fit using conventional indices that were available in both the lavaan and Genomic SEM software (9) (**Supplementary Material 5**). Only data from UKB (the largest sample) was included in the phenotypic factor analyses. For the genetic factor analyses, GWAS summary statistics from the meta-analyses for each AUDIT item were subjected to standard QC, as described above. Genomic SEM’s multivariable version of LDSC was then used to estimate the genetic covariance and sampling covariance matrices (S and V, respectively) for the AUDIT items, which were used to test the specified confirmatory factor models. The S matrix was smoothed beforehand as it was slightly non-positive definite. Factor extension analysis was used to estimate the expected factor loading of item 6, as it was excluded from the final genetic confirmatory factor model due to non-significant SNP heritability.

### Multivariate genome-wide association analyses

We used Genomic SEM (9) v0.0.2 in R to perform multivariate GWAS for the latent genetic factors from the best-fitting model. Individual SNP effects were estimated for the latent genetic factors in each model if they (i) were available in all univariate summary statistics, (ii) had a MAF ≥.5%, and (iii) were present in the 1000 Genomes Phase 3 v5 reference panel. The effective sample size for each latent factor was estimated using the approach described by Mallard and colleagues (16).

### Biological annotation, gene and transcriptome-based association analyses

We performed multiple in-silico analyses to compare the results from each of the AUDIT latent genetic factors. First, we used FUMA (20) v1.2.8 to identify independent SNPs and study their functional consequences, which included ANNOVAR categories, Combined Annotation Dependent Depletion scores, RegulomeDB scores. Second, we performed MAGMA v1.08 (20, 21) competitive gene-set and pathway analyses for each of the AUDIT genetic latent factors. SNPs were mapped to 18,546 protein-coding genes from Ensembl build 85. Gene-sets were obtained from Msigdb v7.0 (“Curated gene sets”, “GO terms”). We also used an extension of this method, Hi-C coupled MAGMA (**H-MAGMA**)(22), to assign non-coding (intergenic and intronic) SNPs to genes based on their chromatin interactions. Exonic and promoter SNPs are assigned to genes based on physical position. We used four Hi-C datasets, which were derived from fetal brain, adult brain, iPSC-derived neurons and and iPSC-derived astrocytes (https://github.com/thewonlab/H-MAGMA). Lastly, we used S-PrediXcan v0.6.2 (23) to predict gene expression levels in 13 brain tissues, and to test whether the predicted gene expression showed divergent correlation patterns with each of the AUDIT latent genetic factors. Pre-computed tissue weights from the Genotype-Tissue Expression (**GTEx** v8) project database (https://www.gtexportal.org/) were used as the reference transcriptome dataset. Further details are provided in the **Supplementary Material 6**.

### Polygenic risk score analyses

#### Prediction of alcohol phenotypes in UK Biobank and COGA

We created polygenic scores (**PRS**), using the PRS-CS “auto” version (24), for the latent genetic AUDIT factors, *Consumption* and *Problems*, and their sum score equivalents AUDIT-C and AUDIT-P, amongst European subjects from two independent samples: (i) an independent subset of unrelated individuals of European ancestry in the UKB who did not fill out the AUDIT, and (ii) a subset of individuals of European ancestry from the Collaborative Study on the Genetics of Alcoholism (**COGA**)(25), which includes probands meeting criteria for alcohol dependence, their family members, and community control families. Using the ‘score’ algorithm in PLINK v1.90, we computed individual-level PRS to predict additional alcohol phenotypes (drinking quantity, drinking frequency, and lifetime AUD diagnosis) measured in UKB and COGA (**Supplementary Material 7**). We tested for associations between PRS of AUDIT phenotypes [the latent genetic AUDIT factors (*Consumption, Problems*) and the sum score equivalents (AUDIT-C, AUDIT-P)] and alcohol phenotypes using linear (quantity and frequency phenotypes) or logistic (AUD) regression models in R v3.6.3. In UKB, we included sex, age at first assessment, Townsend Deprivation Index score (26) and the first ten ancestry PCs as covariates. In COGA, we included age, sex, array type, income, and the first 10 ancestry PCs as fixed effect covariates, with family ID included as a random effect (i.e., allowing the intercept to vary by family).

We aimed to compare the performance of the latent factor-based PRSs (*Consumption* and *Problems* PRS) against the performance of their sum score counterparts (AUDIT-C and AUDIT-P PRS) in predicting different alcohol phenotypes. To this aim, we tested two independent models: (i) cross-dimension PRS models (i.e., *Consumption* and *Problems* included as simultaneous predictors), and (ii) cross-method PRS models (i.e., *Consumption* + AUDIT-C PRS included as simultaneous predictors in a model, and *Problems* + AUDIT-P PRS included as simultaneous predictors in a model). We corrected for the total number of outcome phenotypes across the validation samples using a conservative Bonferroni *p* value = 8.33E-3, since the same PRSs were used as predictors across models (and were correlated with each other).

#### Phenome-wide association study in BioVU

PRS were computed using the PRS-CS method (24) (described above) for each of the 66,915 unrelated genotyped individuals of European ancestry from the Vanderbilt University Medical Center (**BioVU**)(27).

We performed phenome-wide association analyses (**PheWAS**) for *Consumption* and *Problems* using the PheWAS (28) v0.12 package in R. We fitted a logistic regression model to each of 1,335 case/control phenotypes to estimate the odds of each diagnosis given the PRS, after adjustment for sex, median age of the longitudinal electronic health record measurements, and the first 10 PCs. We repeated the PheWAS analyses using AUD diagnoses (phecodes 317, 317.1) as additional covariates. We used the standard Benjamini–Hochberg false discovery rate (**FDR 5%**) correction to account for multiple testing.

## RESULTS

### Phenotypic and genetic analyses reveal a consistent two-factor structure of alcohol consumption and problematic use

Phenotypic and genetic analyses showed that AUDIT items were positively correlated with each other, with moderate to large estimates (**Tables S3-4**) except for item 1 (i.e., frequency of consumption), which generally showed lower correlation estimates. In general, we found that genetic correlations were moderately larger than the phenotypic correlations (mean absolute difference = .198), which was driven by stronger genetic correlations among items 4 through 10 (i.e., the problematic alcohol use phenotypes). Of note, all AUDIT items exhibited significant SNP heritability (**Table S5**), except for item 6 (i.e., ‘eye opener’), perhaps because this item showed low rates of endorsement across all three cohorts (**Table S2**). For this reason, we excluded item 6 from all subsequent analyses, and a factor extension analysis was performed to estimate its expected factor loading in the final model.

We found that a correlated factors model provided the best fit (**Figure 1, Tables S6-7**) to both the genetic and the phenotypic covariance matrices [phenotypic model: (χ^2^(26)=4252.963, CFI=.994, SRMR=.041), genetic model: (χ^2^(26)=142.689, CFI=.982, SRMR=.067)]. That is, the patterns of genetic and phenotypic correlations among the AUDIT items could both be represented by a factor model with two correlated factors: one that captured the covariance among alcohol consumption items (items 1-3, hereafter named “*Consumption*”) and one that captured the covariance among alcohol-related problems (items 4-10, hereafter named “*Problems*”). These two latent factors were highly correlated with each other, phenotypically (*r*_p_=.825, *SE*=.002) and genetically (*r*_g_=.801, *SE*=.037). Nearly all items had large factor loadings across both levels of analyses except item 1, which consistently had markedly smaller factor loadings and larger residual variances.

**Figure 1.**
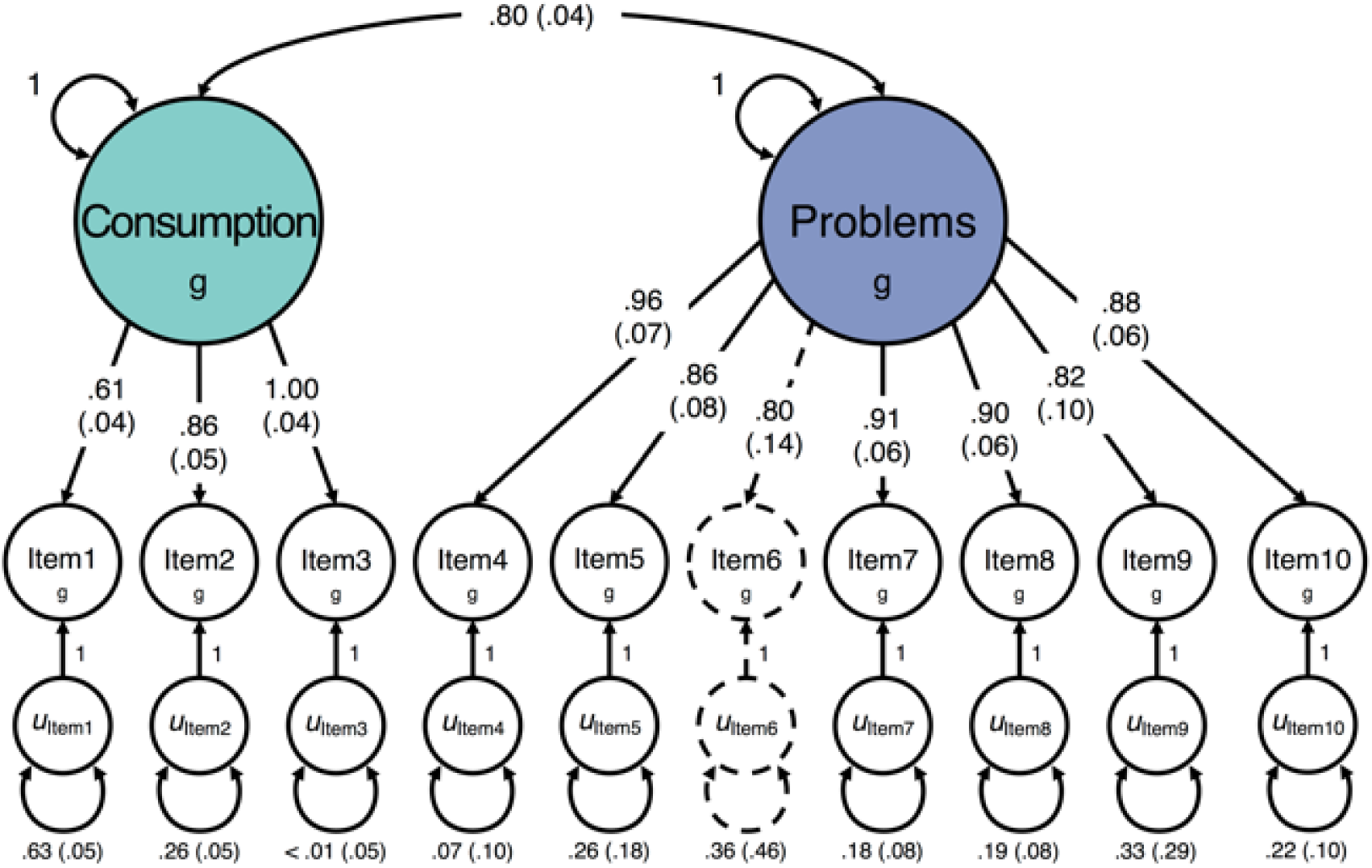
Genetic relationships between AUDIT items. Path diagram of the best fitting genetic confirmatory factor model for AUDIT, as estimated with Genomic SEM. All parameter estimates are standardized, and standard errors are presented in parentheses. The genetic components of items and factors (denoted by g) are inferred variables that are represented as circles. Regression relationships between variables are represented as straight one-headed arrows pointing from the independent variable(s) to the dependent variable(s). Covariance relationships are depicted as curved two-headed arrows linking two variables. The variances for factors are represented as a two-headed arrow connecting the variable to itself, as are the residual variances for individual items (denoted by u). As item 6 was included via factor extension, its parameter estimates are illustrated using dashed lines.

The two correlated-factors model was compared to other solutions. A model with a single common factor provided acceptable fit for the phenotypic (χ^2^(27)=14967.064, CFI=.978, SRMR=.070) and genetic (χ^2^(27)=350.785, CFI=.949, SRMR=.094) factor analyses, but it did not minimize the standardized difference between the observed and predicted correlations as well as the correlated factors model (**Table S7**). The parallel factor model (i.e., the sum score model) exhibited very poor fit, reflected by the strong, unanimous bias observed in the model implied correlations [phenotypic model: (χ^2^(34)=43655.530, CFI=.936, SRMR=.143), genetic model: (χ^2^(43)=607.196, CFI =.911, SRMR=.470)]. Accordingly, we identified the correlated factors model as the best fitting and most appropriate model for further genetic analyses.

### Latent variable approach characterizes and ameliorates bias in GWAS of alcohol consumption

By estimating genetic correlations in a Genomic SEM framework, we identified interesting patterns of relationships between 100 exogenous phenotypes (chosen based on previous findings or hypothesized relationships) and the *Consumption* and *Problems* latent genetic factors. We also examined correlations with the residual genetic variance in item 1 (i.e., the genetic variance in item 1 that is unrelated to other AUDIT items; hereafter named “*Frequency Residual*”). Results are reported in **Table S8**.

For *Consumption* and *Problems*, we found that their patterns of genetic correlation with other phenotypes were much more similar than previously reported for AUDIT-C and AUDIT-P (4). Both *Consumption* and *Problems* showed strong positive genetic correlations with alcohol dependence. *Consumption* and *Problems* were also positively related to other measures of substance use (e.g. cannabis use disorder, impulsivity). Furthermore, the previous positive associations that we observed between AUDIT and indices of socioeconomic status (e.g. educational attainment) were now attenuated.

We did still observe that, compared to *Consumption, Problems* was more strongly related to psychopathology (e.g., post-traumatic stress disorder, depression, bipolar disorder, schizophrenia). We also identified novel divergent associations with pain phenotypes, malnutrition and measures of social satisfaction (e.g., *Problems* showing genetic overlap with these conditions) suggesting that, as we anticipated, the genetic contributions to alcohol consumption and misuse reflect both complementary and distinct genetic factors.

Finally, *Frequency Residual* was negatively associated with alcohol dependence (**Figure 2**). We also found positive genetic correlations between *Frequency Residual* and socioeconomic outcomes, including educational attainment, household income, and intelligence. Furthermore, we observed consistently negative genetic correlations between *Frequency Residual* and other psychiatric and substance use disorders, such as major depressive disorder and cannabis use disorder. This result suggests that many of the puzzling genetic correlations previously reported for alcohol consumption were driven by variance related to socially-stratified differences in behavior rather than variance related to the alcohol phenotypes of clinical interest.

**Figure 2.**
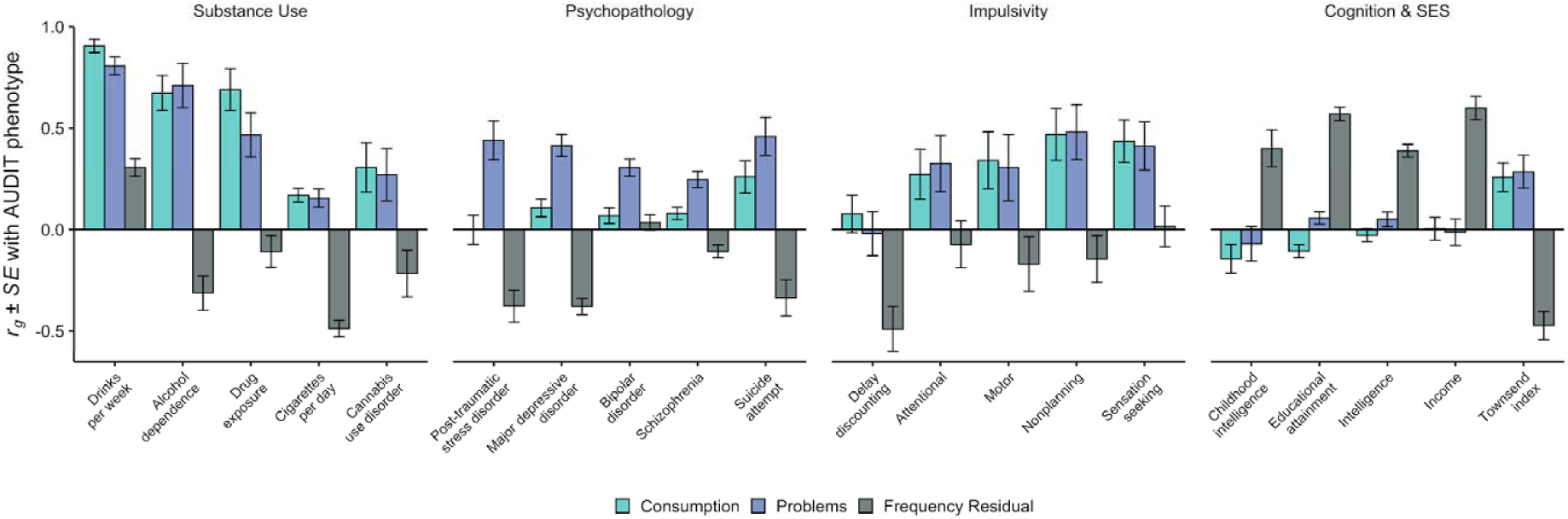
Genetic correlations between latent AUDIT phenotypes and other complex traits. Bar charts of the genetic correlation (*r*_g_) results for three AUDIT phenotypes: *Consumption* (green), *Problems* (blue), and Frequency Residual (gray). Point estimates and corresponding standard errors (*SEs*) are displayed for select phenotypes related to substance use, psychopathology, impulsivity, cognition, and socioeconomic factors. Full results are reported in **Supplementary Table 8**.

### Multivariate GWAS confirm a distinct genetic basis between alcohol consumption and misuse

The results of our multivariate GWAS for *Consumption* and *Problems* are presented in **Figure 3**. We identified 8 independent loci that were associated with *Consumption* (**Table S9**). For *Problems*, we replicated 2 loci on chromosome 4, located in ethanol metabolizing genes (**Table S10**). The signal associated with the latent factors is convergent with that of the sum scores, with a few exceptions (**Supplementary Material 6**.**1**.**1** and **Tables S11-12**).

**Figure 3.**
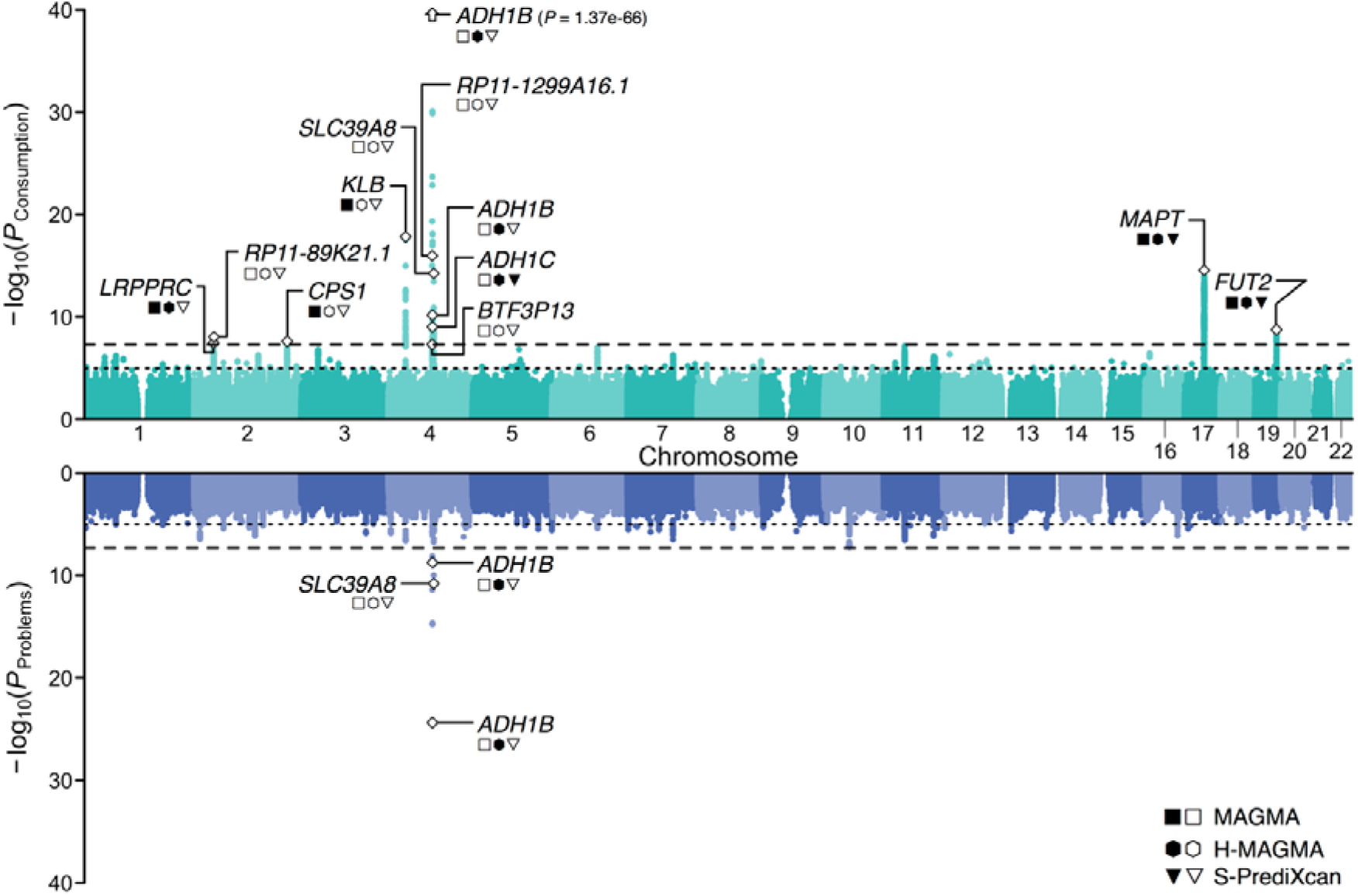
Multivariate genome-wide association analyses for the latent genetic factors. Miami plot for the two latent genetic factors: *Consumption* (top) and *Problems* (bottom). Approximately independent lead SNPs are labeled with a white diamond. For each lead SNP, the neared gene is labeled. Additional symbols convey findings from additional biological annotation; filled symbols indicate that the gene was identified in the corresponding pipeline, while empty symbols indicate that the gene was not. The y-axis refers to the significance on a -log_10_ scale, the x-axis refers to chromosomal position, the horizontal dotted line marks suggestive significance (*p* = 1E-5), and the horizontal dashed line denotes genome-wide significance (*p* = 5E-8).

Some loci included genes that were only associated with *Consumption* (**Table S31**), such as *KLB, RCF1* and the *MAPT/CRHR1* region, which were previously associated with alcohol consumption behaviors (3–5, 29), and other novel candidate genes for alcohol such as *CPS1*, previously associated with metabolic conditions (**Table S13**).

We performed *in-silico* gene-based and transcriptome-based analyses (**Tables S15-30**), which consistently revealed both convergent and divergent associations for *Consumption* and *Problems* (**Table S31**). For example, both factors robustly implicated ethanol metabolizing genes (*ADH1B, ADH1C*) and dopamine transmission [*DRD2*, involved in mediating the rewarding effects of drugs (30)], as well as pleiotropic genes previously implicated in anthropomorphic and metabolic traits [e.g., *CELF1* (5, 31)], and intelligence [e.g., *MTCH2* (32), *FAM180B*/*NDUFS3* (33)].

Lastly, gene-set analyses revealed that genes more closely linked to cellular responses to alcohol drinking (e.g., cellular response to retinoic acid) were associated with *Consumption* (**Table S17**), while the gene-sets related to postsynaptic modulation of chemical synaptic transmission were associated with *Problems* (**Table S18**).

### Polygenic risk analyses

#### UKB

In UKB, we found that both *Consumption* and *Problems* PRSs were robustly associated with drinking frequency, drinking quantity, and lifetime AUD (**Figure 4**). However, *Consumption* PRS outperformed (e.g. higher variance explained) *Problems* PRS for alcohol consumption phenotypes (**Table S32**). When comparing the latent factor-based PRSs to their sum score counterparts, *Consumption* PRS outperformed AUDIT-C PRS in predicting AUD diagnosis and drinking quantity (but not frequency), while AUDIT-P PRS outperformed *Problems* PRS across all three phenotypes (**Table S33**).

**Figure 4.**
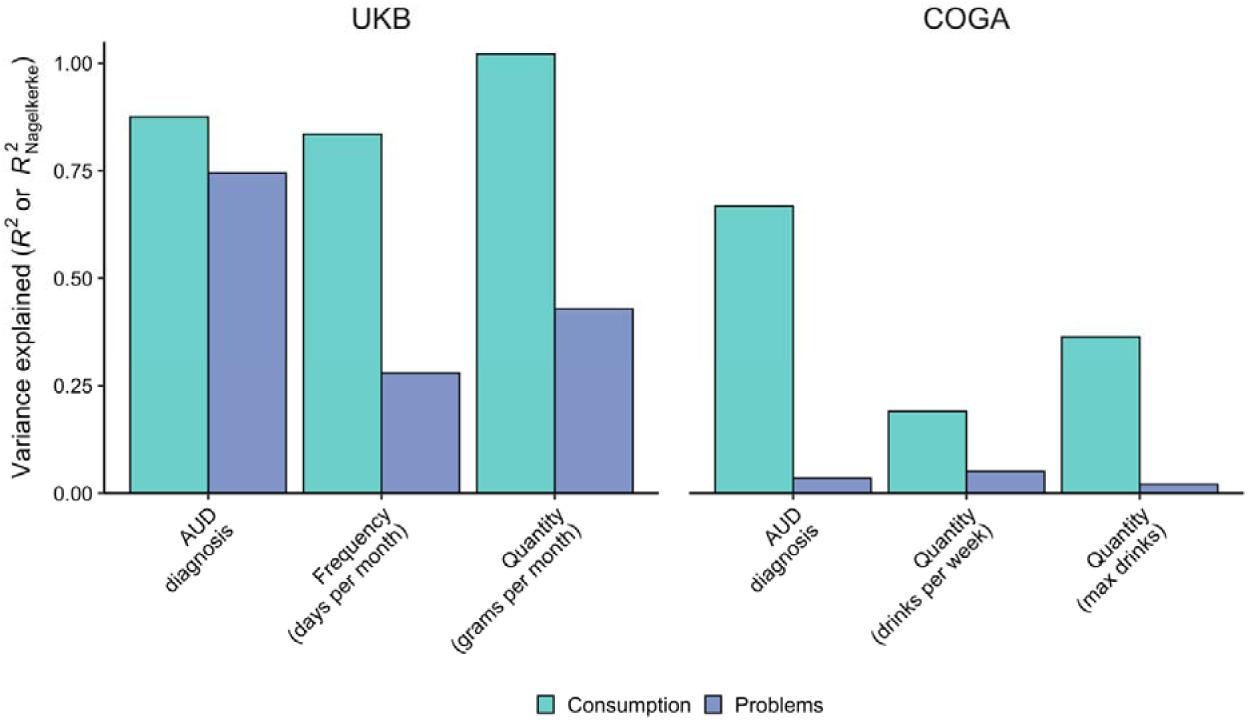
Associations between Consumption and Problems PRS and selected alcohol-related phenotypes. Bar charts of the variance explained by *Consumption* and *Problems* PRS for various clinical and quantitative measures of alcohol use. Values correspond to *R*^2^ or Nagelkerke’s *R*^2^ depending on the use of linear or logistic regression. Results for the independent UKB subset are presented on the left, while results for the independent COGA cohort are presented on the right. Complete results are available in Supplemental Tables 32-35.

#### COGA

In COGA, PRS results were fairly similar to those observed in UKB, with a few exceptions. When both *Consumption* and *Problems* PRS were included in the same model, only *Consumption* PRS showed significant associations with drinks per week, MaxDrinks, and AUD (**Table S34**). When comparing the latent factor-based PRSs to their sum score counterparts, *Consumption* outperformed AUDIT-C PRS, whereas AUDIT-P PRS outperformed *Problems* (**Table S35**). Interestingly, in those models, we found that the strongest associations were between *Consumption* PRS and AUD, and AUDIT-P PRS and AUD.

#### BioVU

We performed two independent PheWAS of *Consumption* and *Problems* PRSs to identify whether these two factors would show different patterns of genetic associations with medical outcomes. Of 1,335 phenotypes, 15 were FDR-significantly associated with *Consumption* (**Figure 5, Table S36**) and 17 with *Problems* (**Table S37**). Both factors were significantly associated with AUD and other tobacco and substance use disorders. Replicating our previous results for AUDIT-C and AUDIT-P, we observed paradoxical negative associations between *Consumption* and metabolic conditions, including diabetes mellitus and obesity phenotypes, whereas *Problems* was primarily positively associated with other psychiatric disorders, including depression, anxiety disorder, bipolar disorder, schizophrenia and suicidal ideation or attempt. Intriguingly, *Problems* was also negatively associated with type 2 diabetes with renal manifestations. Most of the associations did not persist after correcting for AUD, although the direction of effects remained consistent (**Tables S38-39**).

**Figure 5.**
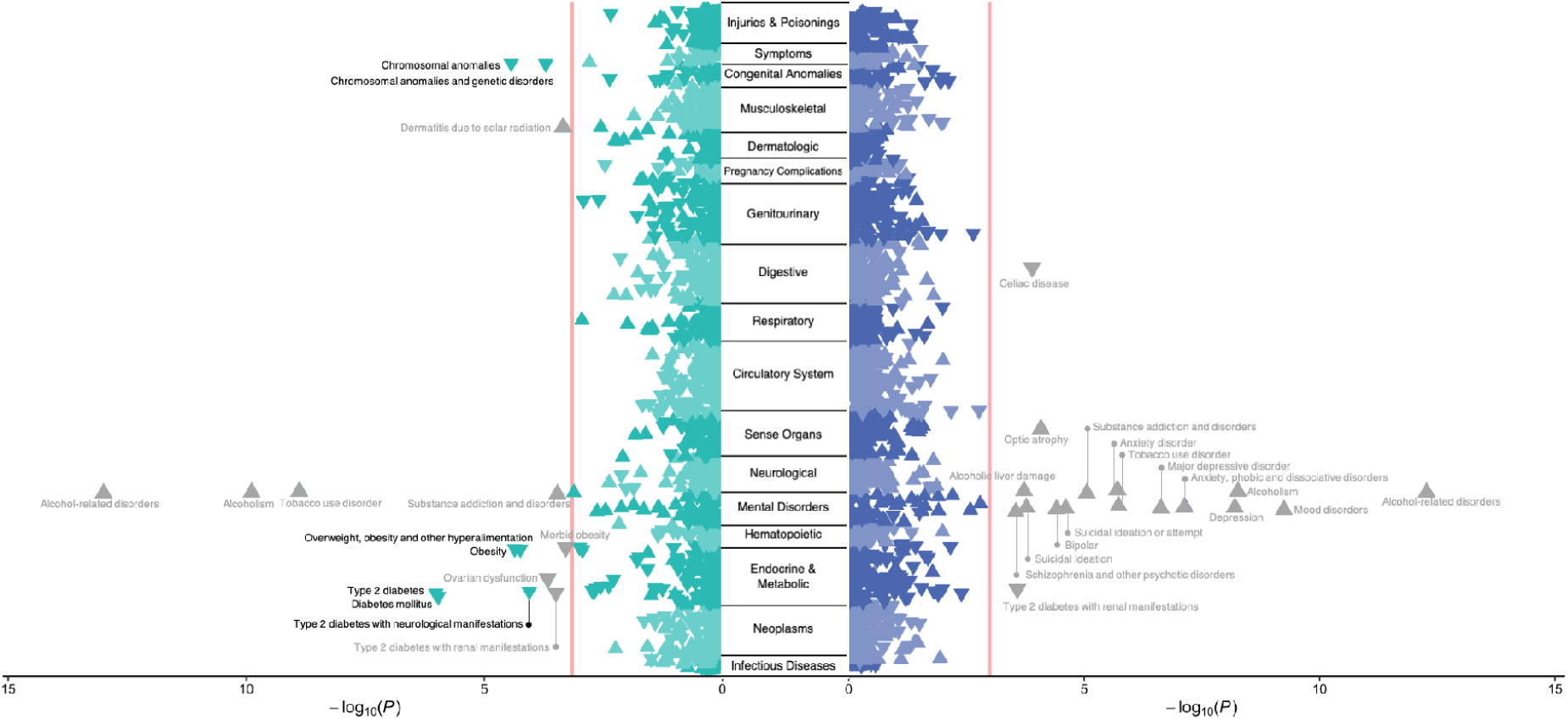
Phenome-wide association study of polygenic risk scores for *Consumption* (left panel) and *Problems* (right panel) against 1,338 diseases available in the biobank from Vanderbilt University Medical Center, BioVU. PheWAS of both *Consumption* and *Problems* revealed positive genetic associations with alcohol use disorders. *Problems* was positively genetically associated with multiple psychiatric conditions, whereas *Consumption* was counterintuitively negatively associated with metabolic conditions. Importantly, most of the associations disappear when adjusting for alcohol use disorder diagnosis (non-significant associations are highlighted in gray).

## DISCUSSION

In the present study, we have performed the first item-level and largest GWAS of AUDIT to date (N=160,824), and used Genomic SEM to elucidate the genetic etiology of alcohol consumption and problematic alcohol use. By conducting phenotypic and genetic factor analyses of the individual AUDIT items, we provide evidence that two correlated latent factors (*Consumption* and *Problems*) parsimoniously explained the covariance in measures of alcohol consumption and problematic alcohol use across both levels of analysis. Moreover, by applying empirically-derived weights to the AUDIT items in a Genomic SEM framework, we demonstrated that our method can ameliorate confounding biases that have complicated previous work with consumption phenotypes (in particular, the bias present in item 1). Notably, both *Consumption* and *Problems* share a strong, positive genetic correlation with alcohol dependence (both *r*_g_∼0.7), and we show, for the first time, that the polygenic signal of *Consumption* is strongly associated with several AUD phenotypes in three independent cohorts. Finally, the results of our bioinformatic analyses further illustrate that *Consumption* and *Problems* have unique components of their genetic etiology. Collectively, our novel framework provides a means to study two genetic liabilities that are more closely related to AUD, and advances our understanding of the associated biology in several ways, as we delineate below.

First, we built upon recent investigations of the genetic etiology of AUDs and related traits by analyzing each of the ten unique items that comprise AUDIT. At this higher resolution, we were able to identify sources of genetic heterogeneity among the items, such as the consistently weaker genetic correlations between frequency of alcohol consumption (item 1) and other drinking patterns (items 2-3) and AUD symptoms (items 4-10). Our item-level approach also allowed us to empirically model the genetic relationships between AUDIT items, providing the first empirical evidence of a correlated two-factor structure for AUD symptoms at the genetic level of analysis. In doing so, we also generated empirically-derived weights to determine how individual items contribute to aggregate measures of alcohol consumption and problematic use. This is an important advance from most quantitative or dimensional genetic studies of AUDs (and other forms of psychopathology), which often use composite score measures that have not been statistically justified.

Second, and perhaps most importantly, we found that *Consumption* was a good genetic proxy of AUD when appropriate weights were applied to the individual items using Genomic SEM. This is a striking change from previous investigations into the divergent genetic bases of alcohol consumption and problematic use, including our own prior analyses of AUDIT. GWASs of alcohol consumption phenotypes have consistently reported low-to-moderate overlap with AUDs that have surprised many researchers (2–5), and even paradoxical negative associations with a variety of diseases and disorders. Our multivariate approach has ameliorated these issues, producing an aggregate measure of alcohol consumption that is more consistent with the known patterns of alcohol phenotype associations established in the existing body of literature, such as a strong genetic correlation with alcohol dependence. Furthermore, we used genetic correlation analyses to characterize the residual genetic variance in frequency of consumption (*Frequency Residual*) that is unrelated to other AUDIT items. These analyses revealed that *Frequency Residual* had consistently positive associations with measures of socioeconomic status and consistently negative associations with measures of substance use and psychopathology. Indeed, these genetic correlations are very similar to those observed in GWASs of AUDIT-C (4, 5) and other GWAS of alcohol consumption (3, 4), suggesting that single-item frequency-based measures of alcohol consumption may be particularly susceptible to measurement error and selection bias. This degree of bias, we speculate, will likely vary from population to population.

Third, we confirmed that the genetic contributions to alcohol consumption are partially distinct from those pertaining to problematic consequences of alcohol use. *In-silico* analyses revealed the value of dissecting the two phenotypes, as gene-and transcriptome-based analyses identified partially divergent biological mechanisms for *Consumption* and *Problems*. For example, the corticotrophin receptor gene (*CRHR1*), which has been associated with alcohol use in animals and humans (34), was associated with *Consumption* only. As a result, we are now beginning to uncover genetic signals for aspects of alcohol involvement that have the potential to be further analyzed at the molecular, cellular and circuit level in cellular and animal model systems.

Fourth, we found that *Consumption* PRS was strongly associated with AUD even in higher-risk cohorts like COGA. This demonstrates the important downstream effects of allowing items to have different weights in phenotype construction. Whereas our current and previous PRS for AUDIT-C have been disproportionately influenced by a single item (frequency of consumption)(35), our *Consumption* PRS was composed of the genetic effects shared among all consumption-focused items. The *Consumption* and *Problems* PRSs were both strongly associated with AUD in UKB – even when both scores were entered in the same model. In COGA, both *Consumption* and *Problems* PRSs were associated with AUD, but *Consumption* PRS was more strongly associated than *Problems* PRS. The increased influence of binge drinking (item 3), which had a large factor loading on *Consumption*, may be partially responsible for these stronger associations in a high-risk sample. However, it is perhaps more likely that these differences might be simply explained by differences in item endorsement and thus predictive power of the discovery GWASs (e.g., *Consumption* had a greater mean χ^2^ than *Problems)*.

Finally, our comprehensive PheWAS analyses have linked different facets of AUD liability (via the latent factor-based *Consumption* and *Problems* PRSs) to a myriad of health-related outcomes in a large, independent biobank. We found that the *Consumption* PRS was consistently negatively associated with a broad range of metabolic and congenital conditions. While it is possible that there is still residual bias in the discovery GWAS, it is important to note that this pattern of paradoxical associations with *Consumption* is not observed in the genetic correlation analyses. Thus, it is possible that these negative associations are illustrative of selection bias or other confounding in BioVU (36), where patients with certain conditions may elect to not drink due to unmeasured factors (e.g., family history, medical advice, contraindications for prescriptions Mirroring the genetic correlation results, we also found that the *Problems* PRS was uniquely associated with numerous psychiatric disorders that are commonly reported to co-occur with AUD. However, and importantly, we identified that the associations between *Problems* PRS and mental health did not persist in the absence of the clinical manifestation of AUD. These findings suggest that the associations with mental health are not the result of horizontal pleiotropy. Instead, they may be either (1) a consequence of AUD, (2) correlated with other risk factors for AUD (along and/or aside from genetic risk), or (3) related to ascertainment of patients with diagnosed AUD in the medical record. These results also encouragingly suggest that treating AUD could have widespread improvements in overall health.

These findings should be interpreted in light of several limitations. Regretfully, AUDIT is a self-report that can be influenced by misreporting, and it only captures alcohol use in the past year, so can be influenced by longitudinal changes in drinking that may be a consequence, for example, of other illnesses (37). People who stopped drinking or never drinkers might represent genetically distinct groups; in our dataset 4,511 individuals were never drinkers, and 4,290 were previous drinkers. While our approach has substantially reduced bias in AUDIT without excluding any individuals from discovery, future studies might consider employing multiple techniques (e.g., separate never drinkers from former drinkers) to further alleviate potential biases associated with frequency of alcohol use in population-based cohorts. Additionally, it remains to be determined how generalizable the genetics of AUDIT are across different populations, especially in samples of different ancestries (as we have only included individuals of European ancestry in the present study) or cultures (UK vs US). A similar point also applies to sex-stratified samples, considering that AUDIT scores differ in men and women. Finally, it is important to note that the *Problems* GWAS is more weakly associated with AUD and other alcohol phenotypes than its AUDIT-P counterpart. We believe this is due to the inclusion of all items with significant SNP *h*^2^, which was critical to the present study – an investigation into the multivariate genetic architecture of AUDIT. However, given our results, it is likely that this loss of power may be attenuated without compromising the factor structure of AUDIT by dropping items with very low SNP *h*^2^ (e.g., item 9).

Analyzing alternative phenotypes as a complementary approach to studying clinically-defined AUD, and psychiatric disorders in general, has generated considerable interest in recent years (38). Collectively, our work demonstrates how AUDIT can inexpensively facilitate such efforts. Here, we have shown that, after correcting for some potential biases, item-or symptom-level analyses can help unpack the genetic etiology of AUD by breaking down genetic influences into specific and shared components; notably, this is only possible because we can contrast our results against gold standard, clinically-ascertained, AUD GWAS datasets. While composite scores have shown some utility in previous genetic association studies, such studies often rely on strong assumptions that the scale is unidimensional, and that each item is equally informative of the construct being measured. In the present paper, we have shown that the latter assumption is false for the AUDIT. In particular, a large proportion of the genetic variance of item 1 appears to be uninformative of a broader consumption construct, as it covaries with. Moreover, although we found a conspicuous degree of unidimensionality among the AUDIT items, our results demonstrate that *Consumption* and *Problems* remain distinct in their associations with human health.

## DATA AVAILABILITY

The GWAS summary statistics for each latent AUDIT factor, and the sum score equivalents (AUDIT-C and AUDIT-P), will be made available on the PGC website.

## ACKNOWLEDGEMENTS

This research was conducted using the UK Biobank Resource under Application Numbers 11425 and 16406.

The AUDIT data collection in ALSPAC was funded by NIH AA018333; ACE was supported by a NIAAA grant (R01AA027522).

The Collaborative Study on the Genetics of Alcoholism (COGA), Principal Investigators B. Porjesz, V. Hesselbrock, T. Foroud; Scientific Director, A. Agrawal; Translational Director, D. Dick, includes eleven different centers: University of Connecticut (V. Hesselbrock); Indiana University (H.J. Edenberg, T. Foroud, J. Nurnberger Jr., Y. Liu); University of Iowa (S. Kuperman, J. Kramer); SUNY Downstate (B. Porjesz, J. Meyers, C. Kamarajan, A. Pandey); Washington University in St. Louis (L. Bierut, J. Rice, K. Bucholz, A. Agrawal); University of California at San Diego (M. Schuckit); Rutgers University (J. Tischfield, A. Brooks, R. Hart); The Children’s Hospital of Philadelphia, University of Pennsylvania (L. Almasy); Virginia Commonwealth University (D. Dick, J. Salvatore); Icahn School of Medicine at Mount Sinai (A. Goate, M. Kapoor, P. Slesinger); and Howard University (D. Scott). Other COGA collaborators include: L. Bauer (University of Connecticut); L. Wetherill, X. Xuei, D. Lai, S. O’Connor, M. Plawecki, S. Lourens (Indiana University); L. Acion (University of Iowa); G. Chan (University of Iowa; University of Connecticut); D.B. Chorlian, J. Zhang, S. Kinreich, G. Pandey (SUNY Downstate); M. Chao (Icahn School of Medicine at Mount Sinai); A. Anokhin, V. McCutcheon, S. Saccone (Washington University); F. Aliev, P. Barr (Virginia Commonwealth University); H. Chin and A. Parsian are the NIAAA Staff Collaborators. We continue to be inspired by our memories of Henri Begleiter and Theodore Reich, founding PI and Co-PI of COGA, and also owe a debt of gratitude to other past organizers of COGA, including Ting-Kai Li, P. Michael Conneally, Raymond Crowe, and Wendy Reich, for their critical contributions. This national collaborative study is supported by NIH Grant U10AA008401 from the National Institute on Alcohol Abuse and Alcoholism (NIAAA) and the National Institute on Drug Abuse (NIDA). ECJ was supported by funding from NIAAA (F32AA027435); AA was supported by funding from K02 DA32573, MH109532, U10AA008401 grants; HJE and JRK were supported by NIAAA U10AA008401.

For the Netherland Twin Register, funding was obtained from the Netherlands Organization for Scientific Research (NWO) and The Netherlands Organisation for Health Research and Development (ZonMW) grants 904-61-090, 985-10-002, 912-10-020, 904-61-193,480-04-004, 463-06-001, 451-04-034, 400-05-717, Addiction-31160008, 016-115-035, 481-08-011, 400-07-080, 056-32-010, Middelgroot-911-09-032, OCW_NWO Gravity program –024.001.003, NWO-Groot 480-15-001/674, Center for Medical Systems Biology (CSMB, NWO Genomics), NBIC/BioAssist/RK(2008.024), Biobanking and Biomolecular Resources Research Infrastructure (BBMRI –NL, 184.021.007 and 184.033.111), X-Omics 184-034-019; Spinozapremie (NWO-56-464-14192), KNAW Academy Professor Award (PAH/6635) and University Research Fellow grant (URF) to DIB; Amsterdam Public Health research institute (former EMGO+), Neuroscience Amsterdam research institute (former NCA); the European Community’s Fifth and Seventh Framework Program (FP5-LIFE QUALITY-CT-2002-2006, FP7-HEALTH-F4-2007-2013, grant 01254: GenomEUtwin, grant 01413: ENGAGE and grant 602768: ACTION); the European Research Council (ERC Starting 284167, ERC Consolidator 771057, ERC Advanced 230374), Rutgers University Cell and DNA Repository (NIMH U24 MH068457-06), the National Institutes of Health (NIH, R01D0042157-01A1, R01MH58799-03, MH081802, DA018673, R01 DK092127-04, Grand Opportunity grants 1RC2 MH089951, and 1RC2 MH089995); the Avera Institute for Human Genetics, Sioux Falls, South Dakota (USA). Part of the genotyping and analyses were funded by the Genetic Association Information Network (GAIN) of the Foundation for the National Institutes of Health. Computing was supported by NWO through grant 2018/EW/00408559, BiG Grid, the Dutch e-Science Grid and SURFSARA. MGN is supported by R01MH120219, ZonMW grants 849200011 and 531003014 from The Netherlands Organisation for Health Research and Development, a VENI grant awarded by NWO (VI.Veni.191G.030) and is a Jacobs Foundation Fellow.

PRS analyses using UKB data were carried out on the Genetic Cluster Computer hosted by the Dutch National computing and Networking Services SurfSARA. DP was supported by NWO VICI 453-14-005.

The dataset(s) used for the PheWAS analyses described were obtained from Vanderbilt University Medical Center’s BioVU which is supported by numerous sources: institutional funding, private agencies, and federal grants. These include the NIH funded Shared Instrumentation Grant S10RR025141; and CTSA grants UL1TR002243, UL1TR000445, and UL1RR024975. Genomic data are also supported by investigator-led projects that include U01HG004798, R01NS032830, RC2GM092618, P50GM115305, U01HG006378, U19HL065962, R01HD074711; and additional funding sources listed at https://victr.vumc.org/biovu-funding/. LKD obtained support from 1R01MH113362, 1R01MH118233 and 1R56MH120736.

SSR was supported by a NARSAD Young Investigator Award from the Brain and Behavior Foundation (Grant Number 27676). YH, MVJ, SSR and AAP were supported by funds from the California Tobacco-Related Disease Research Program (TRDRP; Grant Number 28IR-0070 and T29KT0526).

