## Supplemental Material for "Multivariate GWAS elucidates the genetic architecture of alcohol consumption and misuse, corrects biases, and reveals novel associations with disease"

### 2 Cohorts

The present study used six large, genotyped cohorts: three for discovery and three for validation. To be eligible for inclusion, individuals were required to: (i) be primarily of non-Hispanic European ancestry, (ii) have data on all relevant outcomes and covariates, and (iii) have high-quality, imputed genotyping data. All cohorts were also required to apply appropriate individual- and variant-level quality control thresholds. The cohorts that met these criteria for both analytic stages (discovery and validation) are briefly described below in alphabetical order within each stage.

#### 2.1 Discovery cohorts

Discovery analyses were performed in the three cohorts described below. As genotyping and data collection procedures for each cohort have been extensively described elsewhere, we refer the reader to those studies for details. Quality control thresholds for these cohorts are described in **Supplementary Section 4**.

##### 2.1.1 Avon Longitudinal Study of Parents and Children

The Avon Longitudinal Study of Parents and Children (ALSPAC) is a longitudinal study of 14,541 pregnant mothers living in the former county of Avon, United Kingdom (UK). Details on data collection and genotyping have been described in previous publications (1–3). The ALSPAC website also contains details on all available data through a fully searchable data dictionary (<http://www.bristol.ac.uk/alspac/researchers/our-data/>). In the present study, only the offspring with all relevant data were used in the analyses. Accordingly, the maximum sample size used in discovery analyses was 3,582. Using REDCap (4, 5), participants completed questionnaires that included the AUDIT; for the current analyses, we used responses from the age 22 assessment where available, or responses from age 20 if the individual did not participate at age 22.

All participants provided informed consent at the time of recruitment following the recommendations of the ALSPAC Ethics and Law Committee. Ethical approval for the study was obtained from the ALSPAC Ethics and Law Committee, and the Local Research Ethics Committees. Consent for biological samples has been collected in accordance with the Human Tissue Act (2004). We are extremely grateful to all the families who took part in this study, the midwives for their help in recruiting them, and the whole ALSPAC team, which includes interviewers, computer and laboratory technicians, clerical workers, research scientists, volunteers, managers, receptionists and nurses. The UK Medical Research Council and Wellcome (Grant ref: 217065/Z/19/Z) and the University of Bristol provide core support for ALSPAC. This publication is the work of the authors and S.S-R. will serve as guarantors for the contents of this paper. A comprehensive list of grants funding is available on the ALSPAC website (http://www.bristol.ac.uk/alspac/external/documents/grant-acknowledgements.pdf). This research was specifically funded by NIH AA018333.

##### 2.1.2 Netherlands Twin Register

The Netherlands Twin Registry (NTR) is a national register of twins and multiples, as well as their family members. The NTR follows adolescent and adult twins and their relatives over time, collecting data on numerous complex traits. NTR is a representative sample of the Dutch population (6). Details on data collection and genotyping have been described in previous publications (7, 8). In the present study, the maximum sample size for discovery analyses was 9,975.

All participants provided informed consent at the time of recruitment. Study procedures were performed in accordance with the ethical standards of institutional and/or national research committee, as well as the 1964 Helsinki declaration and its later amendments or comparable ethical standards. The NTR studies were approved by the Central Ethics Committee on Research Involving Human Subjects of the VU University Medical Centre, Amsterdam, an Institutional Review Board certified by the U.S. Office of Human Research Protections (IRB number IRB00002991 under Federal-wide Assurance- FWA00017598; IRB/institute codes, NTR 03-180).

##### 2.1.3 UK Biobank - Discovery

The UK Biobank (UKB) is a population-based biomedical study of genetic and environmental influences on human health and wellbeing. Genotyped participants completed questionnaires related to a myriad of complex traits. A subset of participants also completed an online follow-up protocol, where they answered additional surveys related to mental health and substance use. Details on data collection have been described in previous publications (9), and can also be viewed via the Data Showcase (<https://biobank.ctsu.ox.ac.uk/crystal/>). The maximum sample size used for discovery analyses was 147,267.

All participants provided informed consent at the time of recruitment. Participants also provided consent for follow-up via linkage to their electronic health records. As described by Sudlow and colleagues (10) the UKB investigators developed a robust Ethical and Governance Framework to guide their study procedures, and performed all procedures in accordance with the guidelines established by the relevant ethical, institutional, and regulatory bodies.

#### 2.2 Validation cohorts

Validation analyses were performed in the three cohorts described below. Similar to the discovery cohorts, genotyping and data collection procedures for each validation cohort have been extensively described elsewhere, and we refer the reader to those studies for details. However, as there is more variability in the quality control procedures applied across validation cohorts (*i.e.*, there was no single study-wide pipeline), quality control filters are briefly described below.

##### 2.2.1 Collaborative Studies on Genetics of Alcoholism Study

The Collaborative Study on the Genetics of Alcoholism (COGA) was established to examine familial underpinnings of alcohol use disorders and related behaviors (11–13). Alcohol dependent probands were ascertained from inpatient or outpatient treatment facilities across 7 sites in the United States. Family members of the probands were also invited to participate. Community individuals and their family members were selected from a variety of sources and invited to participate. A proportion of the families in COGA are large and have a high density of alcoholism. In the present study, we excluded participants who (i) were not of European ancestry, (ii) elected not to drink due to personal (*e.g.*, religious, cultural, health) reasons, or (iii) did not report being “regular drinkers” (having at least one drink per month for 6 consecutive months). After these exclusions, our maximum analytic sample consisted of 6,390 individuals.

Analyzed variants were imputed using the 1000 Genomes Phase 3 (14) reference panel. Imputed SNPs with information scores < 0.30 or individual genotype probability scores < 0.90 were excluded, as were SNPs that did not pass Hardy-Weinberg equilibrium (HWE p< 1E-6), and SNPs with a minor allele frequency less than 0.05%. Additional details on genotyping procedures and related quality control can be found in previously published studies (15).

The Institutional Review Boards at all sites approved this study and all participants provided informed consent at every assessment.

##### 2.2.2 UK Biobank - Validation

As the UK Biobank was described in **Supplementary Section 1.1.3**, it is not described again here. For the validation analyses, we selected a subset of UKB, consisting of unrelated White British participants who were not included in the discovery stage analyses (*n*_max_ = 239,719).

Analyzed variants were selected from the third release of the UKB imputed genotype data. In the validation cohort, imputed SNPs were subjected to the quality thresholds reported by UKB investigators, but they were also filtered on several additional thresholds: imputation quality score > .90, minor allele frequency > 1E-5, missingness < .05; unique identifiers only, bi-allelic variants only. SNPs were converted to hard-call format (certainty > .90) prior to analysis.

We included a lower number of PCs (10) in the validation analyses than GWAS (40) to avoid overfitting, given the fact that the analyses are performed on a smaller sample and use aggregate genetic predictors instead of individual variants.

##### 2.2.3 Vanderbilt University Medical Center Biobank

The Vanderbilt University Medical Center (BioVU) biobank cohort is been extensively described previously (16, 17). BioVU is one of the largest biobanks in the United States, consisting of electronic health record data from the Vanderbilt University Medical Center on ~250,000 patients spanning 1990 to 2017. A subset of BioVU patients of European descent (*n*_max_ = 91,602) have been genotyped as part of various institutional and investigator-initiated projects on the Illumina MEGAEX platform, which contains more than 2 million markers. Genotyping and Quality control of this sample have been described elsewhere (16, 17).

Analyzed variants were imputed using SHAPEIT(18)/IMPUTE4(9) with the 1000 genomes phase I reference panel, and variants with INFO < .3 were excluded. A subset of SNPs in linkage disequilibrium was used to calculate relatedness and principal components of ancestry using multidimensional scaling in PLINK v1.9 (19). We randomly removed one individual from pairs of highly related individuals (pihat > .1) to avoid spurious results driven by cryptic relatedness, and restricted to a homogenous population of European descent defined by principal components of ancestry to avoid population stratification effects. Variants were removed if allele frequencies differed significantly (P < 5x10^-5^) between any batch. Finally, we filtered multi-allelic and structural variants, converted dosage data to hard genotype calls, and excluded variants with uncertainty > .1 or INFO < .95

### 3 Phenotype construction

The AUDIT consists of 10 self-report items that are scored via 3- or 5-point ordinal scales. Items 1 through 8 are scored on an ordinal scale ranging from 0 to 4, while items 9 and 10 are scored as 0-2-4. The scoring scheme has been described in our previous GWAS of AUDIT composite scores (20). For consistency, the same phenotyping strategy was applied across all three discovery cohorts. However, in UKB, the AUDIT is administered with ‘gating’ or ‘skip’ logic. This is described in the **Supplementary Section 2.1**, where we also describe the steps taken to minimize the influence of this survey structure.

The distribution of AUDIT scores is shown in **Supplementary Table 2**. Of note, samples broadly differed in their general characteristics: UKB participants were generally older than NTR and ALSPAC, and UKB showed higher socioeconomic backgrounds than the general population (9, 10, 21). The ratio of females was similar across the cohorts (61%, 62%, 57% in ALSPAC, NTR, UKB, respectively).

#### 3.1 Multiple imputation via chained equations in UK Biobank

As noted above, the AUDIT was administered with skip logic in UKB, such that only items 1, 9, and 10 were administered to all participants. Items 2 and 3 were administered based on the response to item 1, and items 4 through 8 were subsequently administered based on the responses to items 2 and 3. Unfortunately, this created patterns of missingness across the AUDIT that affected the sample size, power, and covariance between the items. We have previously assumed that missing responses in the AUDIT represented ‘structural zeroes’ and were re-coded as zeroes (*i.e.*, zero imputation) (20). This approach relies heavily on the assumption that the range of valid responses on item B are entirely contingent on item A (*e.g.*, it is logically impossible for someone to binge drink if they never consume alcohol). While straightforward, this *is* still an imputation model, albeit a restrictive and suboptimal model.

In the present paper, we used multiple imputation by chained equations to minimize the impact of missing data on our item-level analyses. Specifically, we used the MICE package in R to employ a more advanced imputation model based on participants’ income, frequency of any alcohol consumption, frequency of alcohol consumption with meals, current alcohol intake versus previous intake 10 years ago, alcohol use disorder diagnosis status (derived from electronic health records), propensity for risk taking, and depression. We imputed item-level responses that were missing due to skip logic, but did not alter participants’ responses that were intentionally left blank. We imputed the dataset 50 times and used the mean imputed value for each item in subsequent item-level association analyses; however, the composite scores (AUDIT-C and AUDIT-P) were constructed and analyzed using zero imputation (to maintain consistency with previous GWAS). For phenotypic analyses, we used a simpler zero imputation approach (i.e., recoding missing values as zeroes), as the lavaan v0.6.5 package in R cannot readily integrate multiple imputation when using the weighted least squares estimator for categorical variables.

### 4 Genome-wide association analyses

We performed genome-wide association analyses in three population-based cohorts. Each cohort is briefly described below.

#### 4.1 Avon Longitudinal Study of Parents and Children

In ALSPAC, we used PLINK v2 to conduct univariate association analyses for each of the ten AUDIT items, AUDIT-C composite score, and AUDIT-P composite score. As noted in **Supplementary Section 1.1.1**, only offspring genotypes were used in the discovery analyses, as the AUDIT was not administered to mothers. Analyses were further restricted to participants of non-Hispanic European ancestries. The association model included covariate for sex, age (20 or 22), and the first ten principal components of ancestry.

Although we applied the same quality control pipeline to all GWAS results in the present study, some quality control was applied centrally by ALSPAC investigators before we obtained the data. Specifically, markers with > 2% missingness were excluded, as were those with MAF < 0.005, INFO < 0.3, and/or HWE *p* < 5E-7. Individuals missing data on > 5% of markers were excluded. We note this for transparency, but also note that we generally applied more stringent filters ourselves, as outlined in **Supplementary Section 4**.

#### 4.2 Netherlands Twin Register

In NTR, we used the fastGWA function of the GCTA software (22) to perform univariate association analyses for each of the ten AUDIT items, AUDIT-C composite score, and AUDIT-P composite score. The mixed model employed by fastGWA included a random effect with a sparse genetic relatedness matrix in order to correct for kinship present in the sample. We included sex, year of birth, genotyping platform, and the first five principal components of ancestry as covariates in the models.

#### 4.3 UK Biobank - Discovery

In UKB, we used BOLT-LMM v2.3.2 (23) to perform univariate association analyses for each of the ten AUDIT items, as well as the AUDIT-C and AUDIT-P composite scores. We selected BOLT-LMM as our preferred software because it employed a linear mixed model that included a genetic variance component, which (i) permitted the analysis of related individuals and (ii) improved control of population stratification. The genetic variance component was estimated using a set of 483,680 directly genotyped, autosomal SNPs that passed the genotype quality-control, had minor allele frequency greater than 0.005, Hardy-Weinberg-Equilibrium (HWE) *p*-value greater than E–16, and light LD-pruning (window size = 50 kb; variant step-size = 5; *r*^2^ = 0.9) (24). Further details for this pipeline have been previously reported elsewhere (9).

We excluded participants (i) that did not self-report their ethnic background as “White”, “White British”, “White Irish”, or “Any other white background”; (ii) whose self-reported sex and genetic sex were incongruent; (iii) that were identified as purported sex chromosome aneuploidy cases; (iv) that did not pass the sample-level quality control thresholds (9); or (v) that were missing data for the required outcome(s) and covariates. Never drinkers and non-current drinkers were included in the analyses.

The BOLT-LMM association model included covariates for sex, birth year, sex-by-birth year effects, genotyping batch (that also effectively indexed genotyping array), as well as the first 40 principal components of ancestry, which we estimated ourselves (see (25) for details). Analyzed variants were selected from the third release of the UKB imputed genotype data.

### 5 Quality control of association results

Similar to our previous work (25), we used EasyQC (26) to apply a series of thorough quality control filters to our association results. For each cohort, we applied the following thresholds to each set of GWAS summary statistics. Filters were applied in the order that they are listed below.

1. We excluded SNPs if either allele corresponded to a value other than “A”, “C”, “G”, or “T”.
2. We excluded SNPs if any of the following statistics were missing: beta, standard error, *p*-value, effect allele frequency, sample size, and imputation quality score (for imputed SNPs).
3. We excluded SNPs if they had nonsensical or impossible values (*e.g.*, negative standard errors, allele frequencies greater than 1 or below 0).
4. We filtered SNPs with minor allele frequencies less than .005.
5. We removed SNPs with an imputation quality score < .90.
6. In the case of duplicated base pair positions (based on GRCh37), we retained the SNP with the largest sample size.

After applying the filters described above, we inspected diagnostic plots to further ensure that results were not prone to systematic error or bias.

7. We inspected a plot of the allele frequencies in our GWAS sample against those in a non-Hispanic European reference sample in order to check for discrepancies in allele frequencies and errors in strand orientation.
8. To identify discrepancies in reported *p*-values, we examined their relationship the reported coefficient estimates and their SEs.
9. We examined a quantile-quantile plot to evaluate whether population stratification had been sufficiently accounted for in the GWAS.

### 6 Genomic structural equation modeling

Genomic Structural Equation Modelling (Genomic SEM) (27) is a novel statistical method for applying SEM to GWAS summary statistics in order to model the joint genetic architecture of complex traits. Genomic SEM is a flexible framework that allows for empirical modeling of multivariate genetic covariance matrices, such as those estimated via linkage disequilibrium (LD) score regression (28). Here, we used Genomic SEM to conduct a series of analyses in order to study the multivariate genetic architecture of AUDIT items. We highlight three specific aims: (i) identify the latent genetic factor(s) structure that best represent the genetic covariance of AUDIT items, (ii) evaluate genetic relationships between the latent genetic factors and exogenous phenotypes, and (iii) perform multivariate GWAS of the latent genetic factors (pending good evidence of an underlying factor structure).

We have extensively described the general background of Genomic SEM in previous publications (25, 27). Accordingly, we do not review such information in the present materials, although we do explain our project-specific applications of Genomic SEM. We suggest that readers interested in the methodology of Genomic SEM review the original study (27) as well as the online tutorial (<https://github.com/MichelNivard/GenomicSEM/wiki>).

#### 6.1 Confirmatory factor analysis

Genomic SEM can be used to conduct confirmatory factor analysis (CFA), where theoretical models are used to parsimoniously explain the genetic covariances among a set of phenotypes. Here, we use CFA to test a series of competing genetic factor models in order to identify the model that best fit the data. As in traditional CFA, good fit means that the specified latent genetic factor structure adequately explains the observed genetic covariances among the phenotypes.

Fortunately, the AUDIT has been the subject of many factor analytic studies at the phenotypic level (**Supplementary Table 1)**. As the phenotypic null hypothesis suggests that the genetic factor structure will largely be similar to the phenotypic factor structure, we based our models on the existing literature and our own phenotypic factor analysis in UKB (the largest sample), as described in the main text. We used unit variance identification was to set the scale of the latent factors.

We assessed model fit using conventional indices in SEM: the model χ^2^ statistic, the Akaike information criterion (AIC), the comparative fit index (CFI), and the standardized root mean square residual (SRMR). As previously described (27), all of these fit indices retain their standard interpretation within Genomic SEM except for the model χ^2^ statistic, which is used as a comparative measure of fit to evaluate competing models (akin to AIC) rather than a measure of statistical significance. CFI values greater than .90 and SRMR values less than .08 were considered reflective of good model fit (29). While models with good fit also traditionally have non-significant model χ^2^ statistics, the test is extremely sensitive in large samples like the ones used in this study. As such, the model χ^2^ statistic is interpreted as a comparative measure of fit to evaluate competing models rather than a measure of statistical significance.

##### 6.1.1 Factor extension

Factor extension is a method for estimating factor loadings for variables that were not included in the original analysis. In the present study, item 6 was excluded from the final analysis on the basis of its non-significant SNP heritability. Nonetheless, we were still interested in its relationship to the *Problems* factor to which it should theoretically related.

To estimate the putative loading for item 6, we employed a straightforward two-step procedure. In the first step, we freely estimated the correlated factors model again excluding item 6. In the second step, we included item 6 and re-estimated the model; however, we fixed all of the factor loadings and residual variances for items 1-5 and 7-9 to be the same across models. Only the parameters for item 6 were freely estimated in this second step. This yielded an unattenuated estimate of the relationship between item 6 and *Problems* without affecting the overall factor structure.

##### 6.1.2 Genetic correlation

As described in our previous work (25), Genomic SEM can also be used to estimate the genetic correlation between a latent factor and an exogenous phenotype that is not included in the factor model. We note that this method is preferable over bivariate LD score regression for reasons that have been previously explained (25). In the present study, we used this method genetic correlations between 100 exogenous phenotypes and the latent genetic factors, *Consumption* and *Problems*, as well as *Frequency Residual* (*i.e.*, the residual genetic variance in item 1).

### 7 Biological annotation

To link our genome-wide association results to putative risk genes and associated biology, we performed a series of biological annotation analyses described below.

#### 7.1 Functional mapping and annotation

We used FUMA v1.3.5e (30) to study the functional consequences of the lead SNPs for Consumption and Problems, which included ANNOVAR categories (*i.e.*, the functional consequence of SNPs on genes), Combined Annotation Dependent Depletion (CADD) scores (*i.e.*, scores > 12.37 are the suggested threshold to classify a SNP as deleterious), RegulomeDB scores (*i.e.*, biological evidence that the SNP is a regulatory element, scores with 1a representing the strongest evidence).

We used FUMA (30) to identify whether the loci associated with *Consumption* and *Problems* (and SNPs in LD, r2 > 0.1) were previously associated (*p* < 1E-5) with other traits from the GWAS Catalog (version e96 2019-05-03). This information is shown in **Supplementary Tables 13-14**).

##### 7.1.1 Supplemental results for AUDIT-C and AUDIT-P

We identified 8 independent loci for AUDIT-C, showing convergence with *Consumption* with a few exceptions: *GCKR* [previously associated with forms of alcohol behavior [*e.g.* (20, 31–34)], physical activity, sleep patterns and metabolic traits [*e.g.* (35–38)], *ORC5* [implicated in acute myeloid leukemia (39), brain volume (40), alcohol consumption (32)] and *GPR139* [previously associated with smoking behavior (41), anthropometric traits (41, 42), age at menarche (37), insomnia (43)] were only associated with AUDIT-C, whereas others, such as *CPS1* [associated with metabolic and anthropometric phenotypes (43–45) and chronic kidney disease (46)], *FUT2* [pleiotropic gene associated with *e.g.,* alcohol behaviors (20, 32), blood protein levels (47), diarrhea (48), serum cancer antigen levels (49), tumor biomarkers (50), vitamin B12 levels (51)] and *BTF3P13* [associated with alcohol consumption (50, 51) and behavioral disinhibition (52)] were only associated with *Consumption*. The regions including the genes *ADH1B*, *KLB*, *MAPT*, *SLC39A8* and *RP11-89K21.1* were common to both *Consumption* and AUDIT-C.

Compared to the three independent loci that we identified for *Problems*, we identified 6 independent loci for AUDIT-P, including a region containing the well-known dopamine receptor gene, *DRD2*. Several of the loci significant in AUDIT-P [*KLB*, replicating previous GWAS of alcohol phenotypes (20, 32–34)], and two novel candidate genes for alcohol, *RP11-478B11.2*, *AC064875.2*) were not associated with *Problems*, with the exception of the main signal on chromosome 4, including the genes *ADH1B* and *SLC39A8*.

#### 7.2 Gene-based, gene-set, and gene-property analyses

We performed competitive gene-based association analyses using the multivariate GWAS for *Consumption* and *Problems* by applying the “Multi-marker Analysis of GenoMic Annotation” (MAGMA v1.07) (30) in FUMA. SNPs were mapped to 18,857 protein-coding genes from Ensembl build 85 based on physical position. This method accounts for LD patterns, using the 1000 Genomes European sample as a reference. We used a Bonferroni-corrected significance threshold (*p* < 2.65E-6, adjusted for testing 18,857 genes).

We also performed a MAGMA (53) gene-set analysis in FUMA. We used 18,116 curated gene sets and Gene Ontology (GO) terms obtained from the Molecular Signatures Database (MSigDB v7.0, https://www.gsea-msigdb.org/gsea/msigdb/index.jsp) (54), to study the relationships between *Consumption* and *Problems* and the biological processes, molecular function and cellular component of the individual gene products. We used a Bonferroni-corrected significance threshold (*p* < 3.23E-6), adjusted for testing 15,485 gene sets).

#### 7.3 Gene-based analysis of cortical chromatin interaction data

We used H-MAGMA (55), an extension of MAGMA v1.08 that uses Hi-C data to assign non-coding (intergenic and intronic) SNPs, to identify trait-associated genes based on their chromatin interactions. Exonic and promoter SNPs were assigned to genes based on physical position. We used four Hi-C datasets derived from adult brain (56), fetal brain (57), and iPSC derived neurons and astrocytes (58) (all available for download: <https://github.com/thewonlab/H-MAGMA>). The results are reported in **Supplementary Tables 21-28**. We used a Bonferroni-corrected significance threshold (*p* < 9.40E-7, *p* < 9.41E-7, *p* < 9.41E-7, *p* < 9.41E-7, for each analysis, respectively).

#### 7.4 Gene-based analysis of cortical transcriptomic data

We used S-PrediXcan v0.6.2 (40) to analyze predicted gene expression levels in multiple brain tissues, and to test whether the gene expression correlated with the genetic liability of Consumption and Problems. As input data, we used the summary statistics for Consumption and Problems, pre-computed tissue weights from the Genotype-Tissue Expression (GTEx, v8) project database (<https://www.gtexportal.org/>) as the reference transcriptome dataset (59), and covariance matrices of the SNPs within each gene model (based on HapMap SNP set; available to download at the PredictDB Data Repository, <http://predictdb.org>) from 13 brain tissues: anterior cingulate cortex, amygdala, caudate basal ganglia, cerebellar hemisphere, cerebellum, cortex, frontal cortex, hippocampus, hypothalamus, nucleus accumbens basal ganglia, putamen basal ganglia, spinal cord and substantia nigra. We used a Bonferroni transcriptome-wide significance threshold of *p* < 3.52E-6 (184,691 gene-tissue pairs).

### 8 Polygenic risk score analyses

To validate our genome-wide association analyses, we performed polygenic risk score analyses in three independent cohorts. The phenotyping procedures in each cohort are briefly described below.

#### 8.1 UK Biobank - Validation

The phenotypes included in the UKB analyses were: (1) drinking quantity (*i.e.*, typical grams of ethanol consumed per month based on reports of monthly or weekly quantities of various types of alcoholic beverages), (2) drinking frequency (*i.e.*, pseudo-continuous number of typical days drinking per month), and (3) lifetime alcohol use disorder (AUD) diagnosis (*i.e.*, ICD10 codes of alcohol use disorders (F10.1, F10.2, F10.3) or alcoholic liver disease (K70) in hospitalization records, death records, or in-person medical interviews). Because of the small number of AUD cases (*n* = 4,463), a random subset of 20,000 control participants was used in the AUD prediction models to improve the imbalanced ratio. We excluded current (n = 10,093) and lifetime non-drinkers (n = 9,323) from polygenic analyses of current consumption measures (quantity/frequency) and lifetime non-drinkers from analyses of AUD.

#### 8.2 Collaborative Studies on Genetics of Alcoholism Study

In COGA, the phenotypes included in the analyses were: (1) number of drinks per week (*i.e.,* the sum of 4 separate measures: average number of beers/wine/liquor/other alcohol consumed per week); (2) maximum drinks in 24 hours; (3) DSM-5 AUD. The measure of maximum drinks in 24 hours was winsorized [anyone with a value $\geq$ 66 (mean + 3SD’s) was set to 65]. The number of drinks per week and maximum drinks in 24 hours were log-transformed before the analysis to account for skew. We included 6,390 individuals who reported being regular drinkers (having at least one drink per month for 6 consecutive months).

#### 8.3 Vanderbilt University Medical Center Biobank

In BioVU, 1,335 phecodes with at least 100 cases were analyzed. We required the presence of at least two International Classification of Disease codes that mapped to a PheWAS disease category (Phecode Map 1.2 (<https://phewascatalog.org/phecodes)> to assign case status (60).
